# Insertion patterns of *P{lacW}* and *P{EP}* artificial transposons on the third chromosome of *Drosophila melanogaster*

**DOI:** 10.1101/074070

**Authors:** Laura I. Popa, Attila C. Ratiu, Alexandru Al. Ecovoiu

**Affiliations:** University of Bucharest, Faculty of Biology, Department of Genetics, Intrarea Portocalelor Street, No. 1-3, 060101 Bucharest, Romania

**Author notes:** Correspondence author: Alexandru Al. Ecovoiu, University of Bucharest, Faculty of Biology, Department of Genetics, Intrarea Portocalelor Street, No. 1-3, 060101 Bucharest, Romania.

## Abstract

Insertional mutagenesis experiments performed on *Drosophila melanogaster* model often relies on induced mobilization of artificial transposons derived from P mobile element. In an attempt to detect transposition preferences, we accomplished a pilot study concerning the insertional patterns of *P{lacW}* and *P{EP}* constructs in the third chromosome of *D. melanogaster.* Our inventory inquiry considered 2177 insertions of *P{lacW}* and 1646 insertions of *P{EP}* available in FB2016_02 release of FlyBase and revealed insertional hotspots and coldspots in 3L and 3R, but also a preference of both artificial transposons to insert in 3R. The general distribution of *P{lacW}* and *P{EP}* insertions appears to be similar but not identical, probably due to differences in size and molecular architecture of these transposons. Our results may have predictive value for experimental design of insertional mutagenesis screenings, but are also expected to contribute to a better understanding of *P* transposon biology.

## Introduction

Transposons are mobile elements implicated in important genetic mechanisms (Jeffares et al., 2006; Linheiro and Bergman, 2012) and the understanding of their insertion pattern may provide important clues regarding their various functions.

In this pilot study, we analyzed the insertion patterns of *P{lacW}* and *P{EP}* artificial transposons (Bier *et al*., 1989; Rorth, 1996) located on the third chromosome of *Drosophila melanogaster*.

In order to achieve our goal, we interrogated the *Transposons, Transgenic Constructs, and Insertions* item archived in FlyBase (http://flybase.org), which provides a comprehensive database containing information regarding the insertion mapping of curated artificial transposons (*insertion_mapping_fb_2016_03.tsv.gz*). The unzipped .*tsv* file includes data about the genomic and/or cytogenetic coordinates of various types of artificial transposons, including those derived from natural transposons such as *gypsy* or *P.*

The processing of the data file and the selection of the *P{lacW}* and *P{EP}* transposon insertions were performed on Bio-Linux 8 (Field et al., 2006). Nucleotide-level mapped *P* insertions located in the third chromosome were set apart by using grep command with the syntax: *grep “P” insertion_mapping_fb_2016_03.tsv > All_P.tsv*. The newly generated *.tsv* file was the object of successive grep commands, thus generating distinct *.tsv* files for *P{lacW}* and *P{EP}* insertions. The genomic coordinates of the insertions were further used to estimate the cytogenetic localization (chromosomal division).

For a series of insertions indexed in *insertion_mapping_fb_2016_03.tsv* only the cytogenetic region is provided, hence they cannot be selected using grep command. As a consequence, we considered the *.tsv* file and applied a series of filters with *LibreOffice Calc 4.0* and selected only the *P{lacW}* and *P{EP}* insertions located between the 61 and 100 chromosomal divisions. Finally, the data selected by both afore-described methods were manually grouped in eight tables comprising the relevant information needed to perform our analysis (Supplementary files 1 and 2). We considered as insertional hotspots those genes affected by at least 10 insertions but in order to avoid biases regarding the distribution of insertional events, only 10 hits for each of the respective genes were graphically represented (a raw data normalization). Since our analysis was mostly done in a non-automatic manner in this preliminary inquiry, some insertions were overlooked or misplaced, an aspect to be further improved.

## Results and discussions

For this study, a total number of 3823 transposons were analyzed, namely 2177 insertions of *P{lacW}* and 1646 insertions of *P{EP}*.

The chart depicted in Figure 1 reveals the insertion patterns of *P{lacW}* transposon on the third chromosome of *D. melanogaster*.

**Figure 1.**
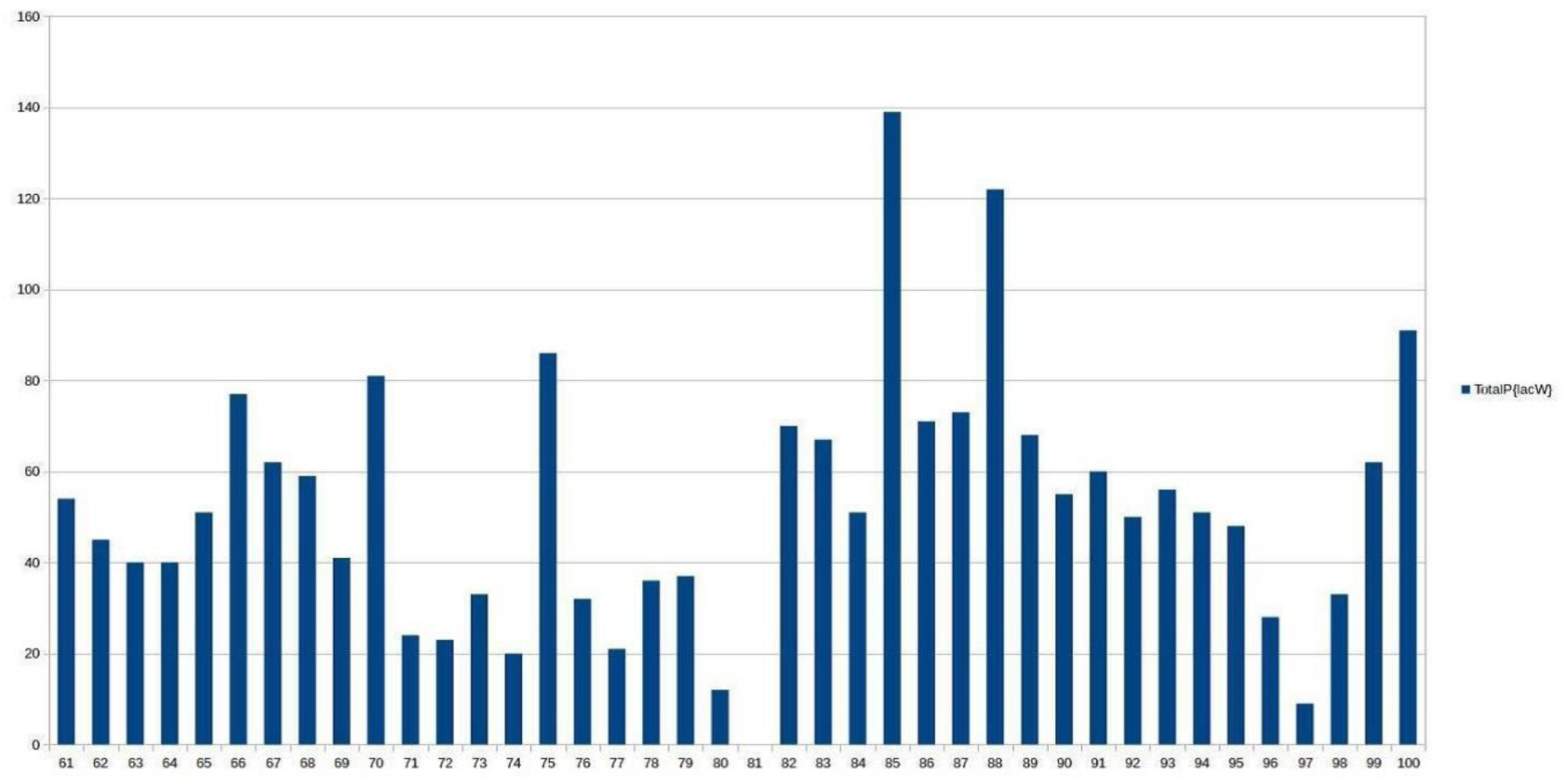
Insertion pattern of *P{lacW}* transposons on the third chromosome of *D. melanogaster.* The x-axis represents the divisions of the third chromosome and the y-axis indicates the number of insertions located on a certain division.

It may be noticed that there is a difference between distributions of *P{lacW}* insertions located on 3L versus those placed on 3R, as 3L harbor a mean of 43.7 insertions/division, while the mean of insertions/division on 3R is 60.2. These sets of data were analyzed with Student’s t-test and the calculated *p* value is statistically significant both after (*p* = 0.0316) and before (*p* = 0.0252) data normalization (statistics were performed using GraphPad Prism 5.03 software).

The lowest number of insertions is registered in the proximity of the centromeric area (for example, the division 81 has no reported insertion). Hotspot divisions are 66, 70 and 75 in 3L and 85, 88 and 100 in the 3R. In contrast, the coldspot divisions are 74, 76 and 80 in 3L and 81, 96 and 97 in 3R (Table 1).

**Table 1.** The hotspot and coldspot divisions in regard to the insertion density of *P{lacW}* on the third chromosome of *D. melanogaster*.

|  | 3L | 3R |
| --- | --- | --- |
| **Hotspot divisions** | 66 (77 insertions) | 85 (139 insertions) |
|  | 70 (81 insertions) | 88 (122 insertions) |
|  | 75 (86 insertions) | 100 (91 insertions) |
| **Coldspot divisions** | 74 (20 insertions) | 81 (0 insertions) |
|  | 76 (32 insertions) | 96 (28 insertions) |
|  | 80 (12 insertions) | 97 (9 insertions) |

A distinct chart describing the pattern of insertions of *P{EP}* transposon on the third chromosome of *D. melanogaster* is depicted in Figure 2.

**Figure 2.**
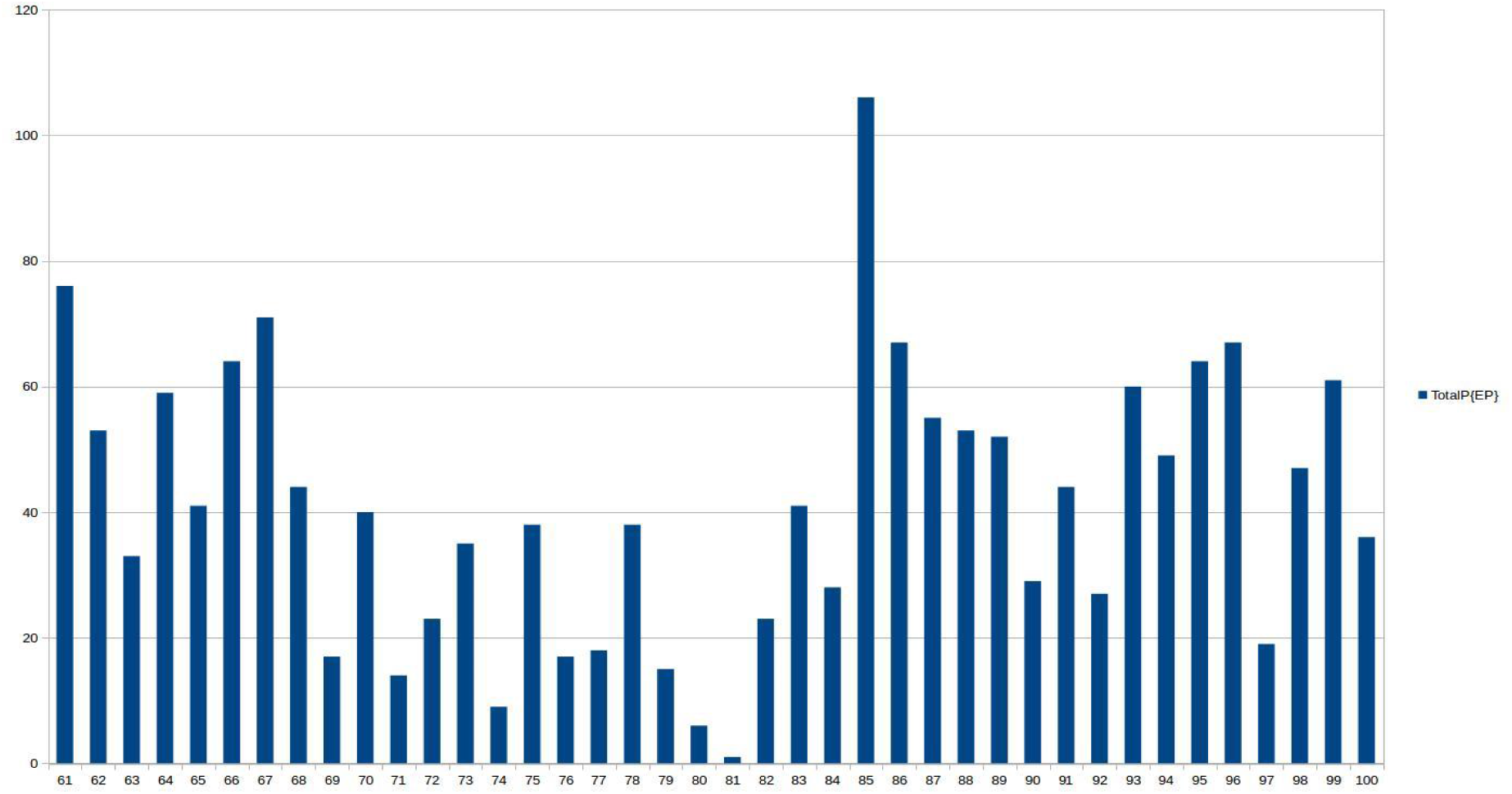
Insertion pattern of *P{EP}* transposon on the third chromosome of *D. melanogaster*. The x-axis represents the divisions of the third chromosome and the y-axis illustrates the number of insertions located in a certain division.

The analysis of the *P{EP}* distribution revealed the same insertional trend as the one observed for *P{lacW}*, with a mean of 35.55 insertions/division for 3L and a mean of 46.45 insertions/division for 3R. Student’s *t test* was applied to these data and the resulting *p* value is not statistically significant after data normalization (*p* = 0.0612), in contrast to the *p* value obtained before the data normalization (*p* = 0.0383). This could indicate that a higher number of hotspot genes contain more than 10 *P{EP}* insertions as comparative to the *P{lacW}* hotspot genes, or the number of *P{EP}* located on specific hotspots tends to be greater than that of *P{lacW}* within their respective hotspots.

The lowest number of insertions were registered in the proximity of the centromeric area (division 81 has only one reported insertion). Hotspot divisions are 61, 66 and 67 in 3L and 85, 86 and 96 in 3R, while the coldspot divisions are 71, 74 and 80 in 3L and 81, 82 and 97 in 3R (Table 2).

**Table 2.**
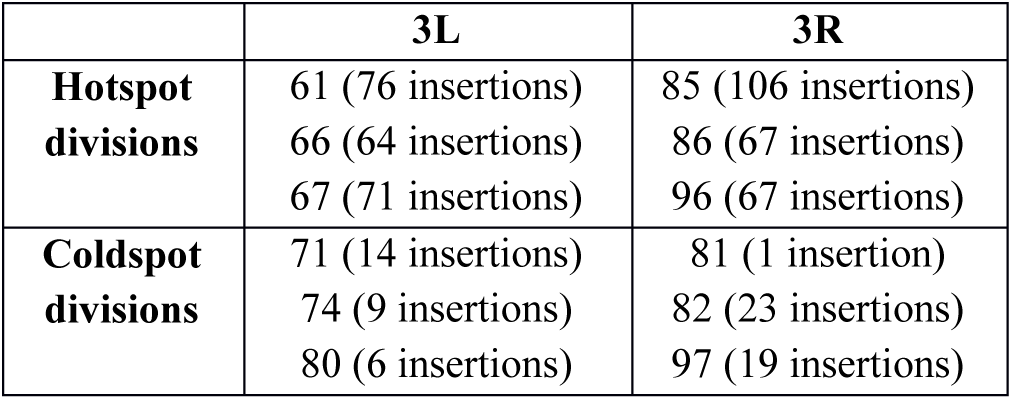
The hotspot and coldspot divisions of *P{EP}* artificial transposon within the third chromosome of *D. melanogaster*.

A comparison of spatial and numeric distribution patterns of both *P{lacW}* and *P{EP}* mobile elements is displayed in Figure 3.

**Figure 3.**
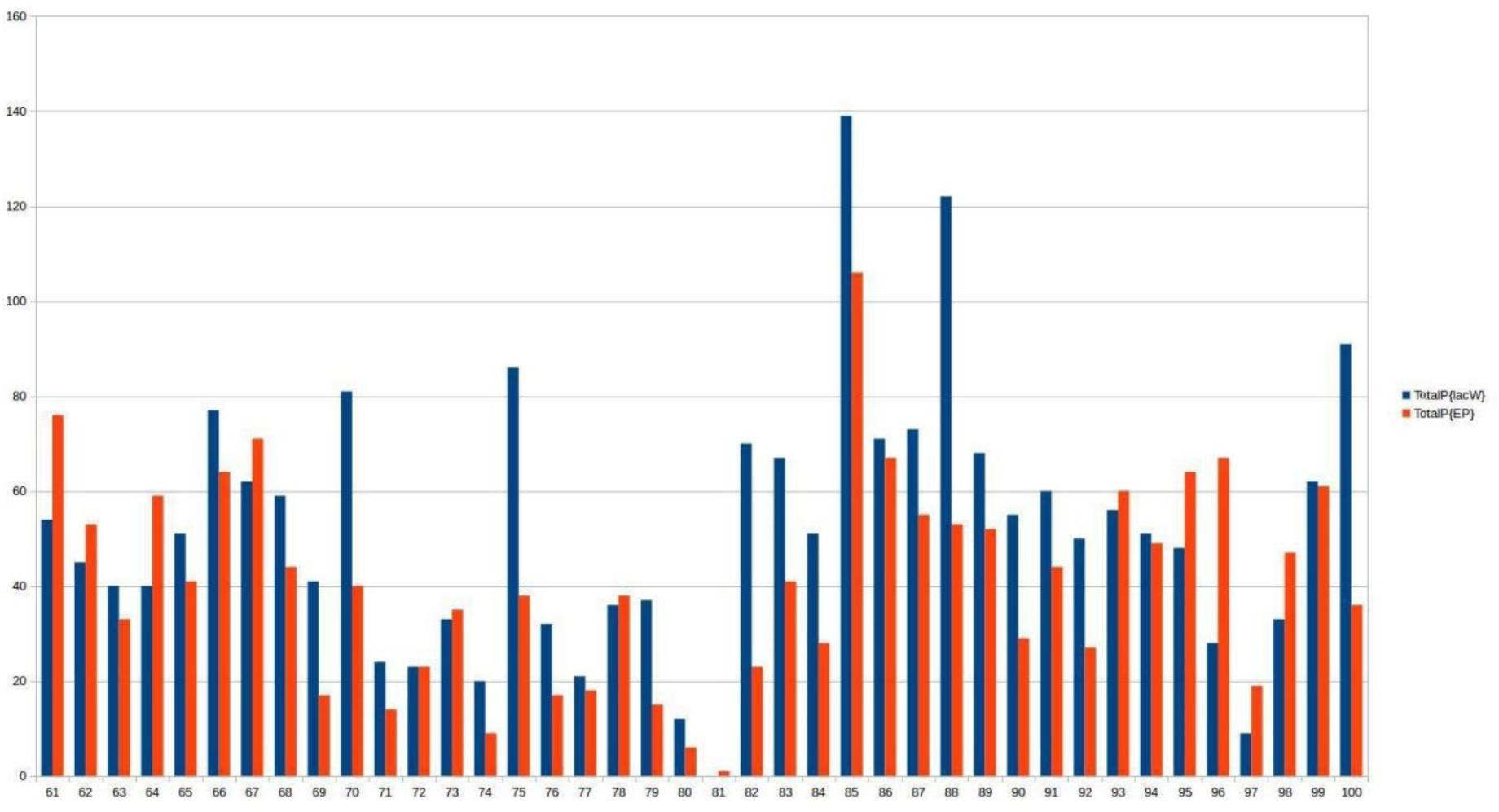
Juxtaposed insertion patterns of *P{lacW}* and *P{EP}* on the third chromosome of *D. melanogaster.* The x-axis shows the third chromosome divisions and on the y-axis there are represented the number of insertions located on each division.

The comparative approach reveals a common tendency of both *P{lacW}* and *P{EP}* artificial transposons to insert on certain divisions of the third chromosome of *D. melanogaster*. The two mobile elements score the lowest number of insertions on division 81 (none for *P{lacW}* and one for *P{EP}*) and the highest number within division 85 (139 for *P{lacW}* and 106 for *P{EP}*). There are also several differences regarding the distribution and richness of insertional hotspots and coldspots, which could be partially explained by the lower number of *P{EP}* transposons included in this study and by the structural differences between the two types of transposons.

A similar study regarding the distribution of *P{lacW}* insertions on the third chromosome offers similar results (Deak *et al*., 1997). As in our analysis, the insertional hotspots on 3L are divisions 66, 70 and 75 and the coldspot division is 80. The pattern of *P{lacW}* insertions affecting 3R is also very similar to the one obtained in our study, with the hotspots located on the same divisions (85, 88 and 100). The lowest number of insertions (coldspots) are also harbored by divisions 81, 97 and 96.

The high degree of similarity between Deak’s study and our work is explained by the fact that a substantial part of data concerning the insertions of *P{lacW}* element, as provided by FlyBase, were obtained during the experiments reported by Deak (around 1200 out of 1813 insertions mapped at cytogenetic level). It is interesting to note that the insertion pattern was not affected by the about 600 insertions reported in FlyBase since the work of Deak *et al*.

## Conclusions

The insertion patterns of both *P{lacW}* and *P{EP}* artificial transposons are significantly different between the two arms of the third chromosome of *D. melanogaster*, with a noticeable tendency towards the insertion in 3R. The hotspot and coldspot divisions are distributed following a similar trend regardless of the considered artificial transposon.

The inquiry methodology used in our pilot study is to be applied for all of the chromosomes of *D. melanogaster* in order to have a broader view of the insertional patterns. Our results may be useful when designing insertional mutagenesis experiments, but also for a deeper understanding of transposon biology.

